# CIDP: A multi-functional platform for designing CRISPR sgRNAs

**DOI:** 10.1101/2022.09.07.506696

**Authors:** Dong Xu, Jin Zhang, Xianjia Zhao, Heling Jiang, Xiongfeng Ma, Weihua Pan

**Author notes:** These authors contributed equally to this work (Dong Xu, Jin Zhang, Xianjia Zhao).

## Abstract

Most of sgRNA-design tools can be run under the precondition of the choice of closely-related species. However, it is hard to select an ideal closely-related species, as more and more different species were studied, and this situation was particular seriously in plant studies. Here, we introduced a new software, CRISPR Integrated Design Platform (CIDP), to solve the problem by allowing users to input genomic sequences for designing sgRNAs. The main function of CIDP was to design sgRNAs after building the sgRNA database using the input genomic sequences. Furthermore, in order to minimize the off-target effects, CIDP will search sgRNAs that appear only once across the whole genome on the target sequences. Meanwhile, CIDP set relevant functions to identify shared sgRNAs of a group of genes. Moreover, we also set primer design and sequence extraction functions in CIDP to help users design sgRNAs efficiently.

---

CRISPR (Clustered Regularly Interspaced Short Palindromic Repeats) technological system has been widely applied in altering phenotypes of various organisms by modifying gene expression. Changing DNA sequences of target genes is one of the most outstanding characters of CRISPR system distinguished with other technologies (Zamore et al., 2000), whose precondition is the accurate binding of single-guide RNA (sgRNA) to the genes of interest (Manghwar et al., 2020). Therefore, sgRNA is the crucial factor influencing the success of CRISPR system. Similar sequences in the global genome with the sgRNA will seriously affect the binding between sgRNA and target genes. How to design sgRNA to minimize off-target cleavage is a great challenge in using CRISPR system (Yan et al., 2018). This is the reason why lots of sgRNA-design tools will calculate off-target scores (Table S1). Most of sgRNA-design tools collected genomes of different species as possible as they can, as background datasets for off-target effects. However, even though the closely-related species were contained in the database of tools, there may also huge differences in genome sequences between the species of interest and the selected closely-related species, which may lead to design errors. Selecting unique sequences in the genome as the sgRNA is an effective way to reduce the off-target effects (Cho et al., 2014). Whereas, few tools can be used for screening unique segments as the sgRNAs.

Here, we introduced a new software, CRISPR Integrated Design Platform (CIDP), which allowed users to build the required database using genomic data of the interested species (Figure 1A). Furthermore, CIDP provides 20 PAM (protospacer-adjacent motif) models for users to select. Meanwhile, the visualization and primer design functions were also set in this software. Designing common sgRNAs can effectively knockout several genes at the same time (Bewg et al., 2022) to produce the required mutants. Therefore, we set related functions to search the shared sgRNAs for the input genes. Considering that users may need to design sgRNAs utilizing the sequences of different parts of interested genes (such as promoter sequence). Functions in “other functions” module of CIDP can be performed to extract the relevant sequences from genomes (Figure 1B). The third module was set for designing the primers for sgRNAs (Figure 1C)

**Figure 1.**
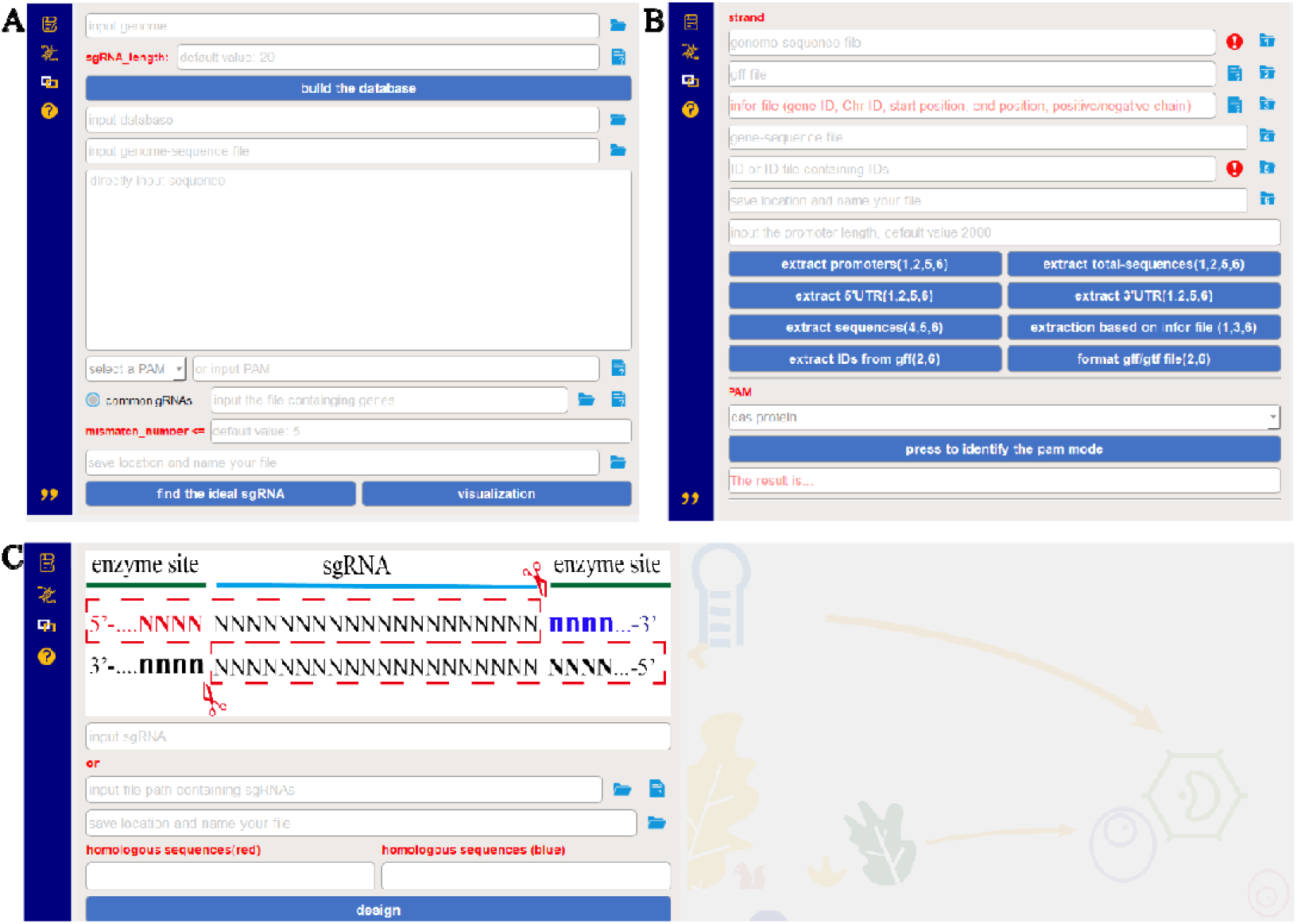
Functional modules in CIDP.

The workflow of CIDP was shown in Figure 2. The unique-sgRNA database was built by the genomic sequences of interest, which contained unique sgRNAs across the whole genome. According to the selected PAM model, CIDP will search all the candidate segments on the sense and antisense chains. Then, the candidate segments will be aligned to the unique-sgRNA database for identifying the unique sgRNAs among of the candidate segments. Furthermore, CIDP will align the obtained unique sgRNAs to the genomic sequences for finding the possible mismatch sites and calculating the off-target scores. Moreover, the hairpin structures and Tm values, as important factors affecting the binding of sgRNAs and target genes, will be provided simultaneously. After that, users can visualize the obtained results and design the relevant primers. The sequence-extraction and format-conversion functions were set in “other functions” module. Users can utilize the extraction function to obtain the related sequences of the interested genes, such as promoters, 5’UTR (Untranslated Regions), 3’UTR, total sequences. The extraction functions were performed based on the information of the input GFF (Generic Feature Format) file. However, some users may not have the GFF files. Therefore, we set the “format conversion” function for transforming GTF (Gene Transfer Format) files to GFF files. Moreover, if users did not have the GFF or GTF files, CIDP can also allow users to input gene-position files for sequence extraction.

**Figure 2.**
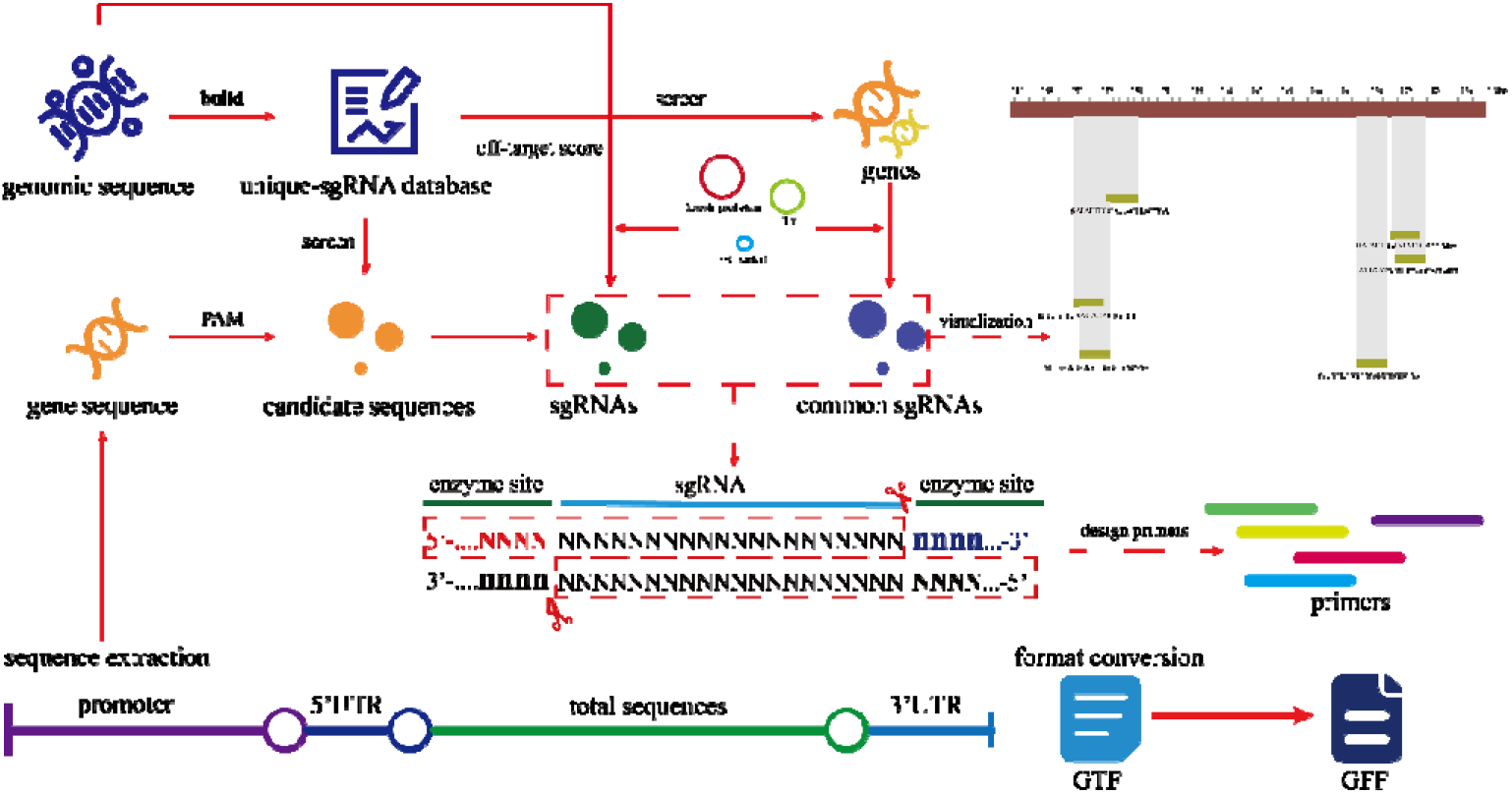
The workflow of CIDP.

The designing results were shown in Figure 3. The designing sgRNAs will be displayed in the right table. Apart from the sequences of sgRNAs, eight parameters about the sgRNAs will also be shown at the same time, including the genomic location, sense strand or antisense strand, GC content, hairpin structures, Tm value, the activities of sgRNAs, the number of off-target sites and off-target scores (Figure 3A). If the exclamation mark was followed in the sgRNA sequences, it is indicated that multiple consecutive adenine (A) or thymine (T) were detected in the sequences of sgRNAs (Figure 3B, the red box). The sgRNAs with sequence uniqueness will be preferentially output. Otherwise, the sgRNAs will be marked by “not_unique” (Figure 3C, the red box).

**Figure 3.**
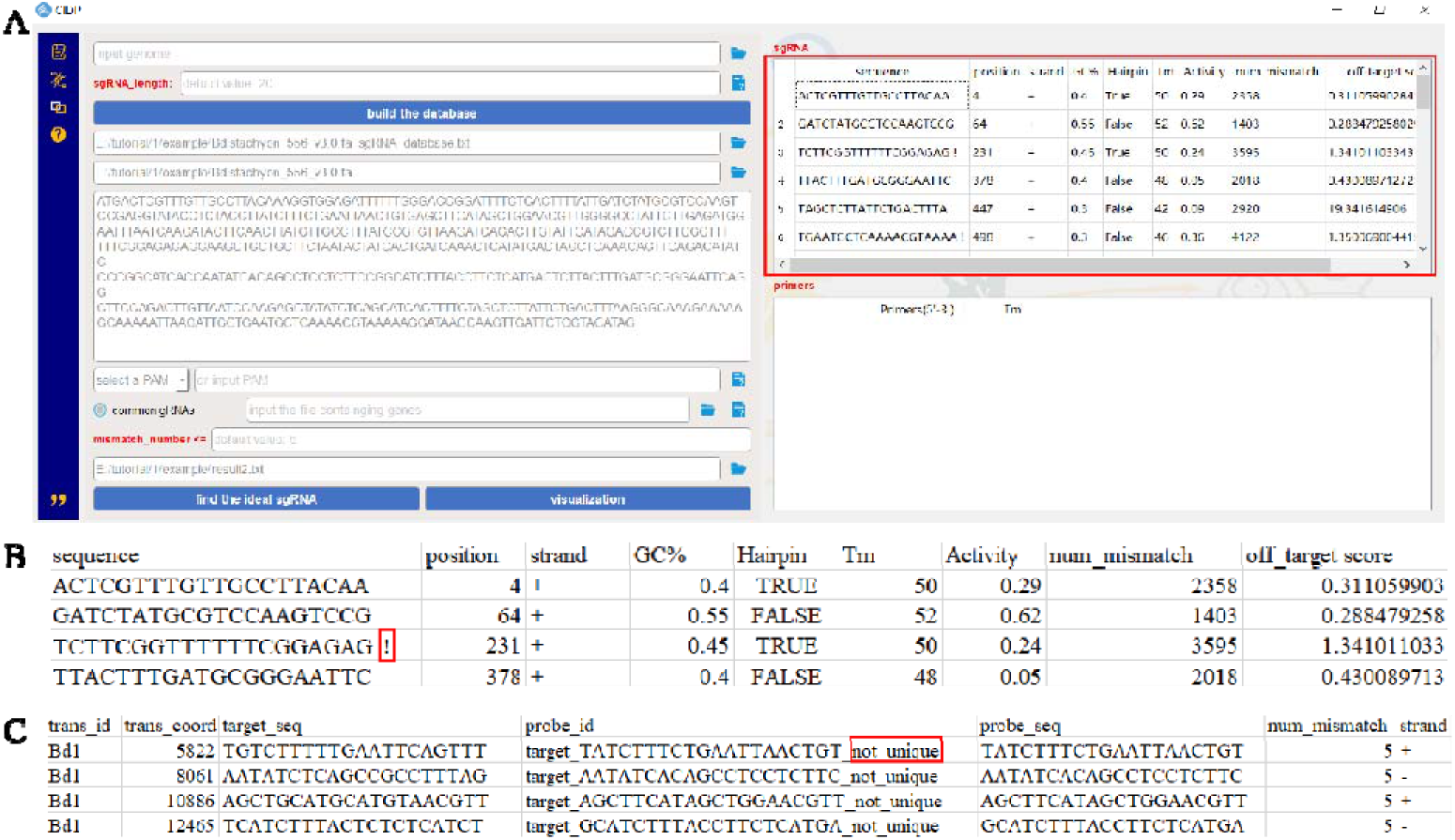
the output results of CIDP.

## SUMMARY

CIDP can perform the sgRNA-design functions, which allows users to build the related database using genomic sequences instead of selecting a closely-related species as the background dataset. Furthermore, CIDP will preferentially output the sgRNAs whose sequences are unique across the whole genome for minimize the off-target effects.

## AVAILABILITY

CIDP was developed by python3.8 with a user-friendly interface. All operations can be done without entering any commands.

## AUTHOR CONTRIBUTIONS

W.P., J.Z. and X.M. conceived the study and designed the experiments. D.X. and X.Z. developed the CIDP software. D.X., H.J. and W.P. wrote the manuscript. X.M. and J.Z. revised the manuscript. All authors approved the final version.

## Reference

Bewg WP, Harding SA, Engle NL, Vaidya BN, Zhou R, Reeves J, Horn TW, Joshee N, Jenkins JW, Shu S (2022) Multiplex knockout of trichome-regulating MYB duplicates in hybrid poplar using a single gRNA. Plant Physiology 189: 516–526

Cho SW, Kim S, Kim Y, Kweon J, Kim HS, Bae S, Kim J-S (2014) Analysis of off-target effects of CRISPR/Cas-derived RNA-guided endonucleases and nickases. Genome research 24: 132–141

Manghwar H, Li B, Ding X, Hussain A, Lindsey K, Zhang X, Jin S (2020) CRISPR/Cas systems in genome editing: methodologies and tools for sgRNA design, off-target evaluation, and strategies to mitigate off-target effects. Advanced science 7: 1902312

Yan J, Chuai G, Zhou C, Zhu C, Yang J, Zhang C, Gu F, Xu H, Wei J, Liu Q (2018) Benchmarking CRISPR on-target sgRNA design. Briefings in Bioinformatics 19: 721–724

Zamore PD, Tuschl T, Sharp PA, Bartel DP (2000) RNAi: Double-Stranded RNA Directs the ATP-Dependent Cleavage of mRNA at 21 to 23 Nucleotide Intervals. Cell 101: 25–33

